## Extended Data for "Detection of isoforms and genomic alterations by high-throughput full-length single-cell RNA sequencing in ovarian cancer"

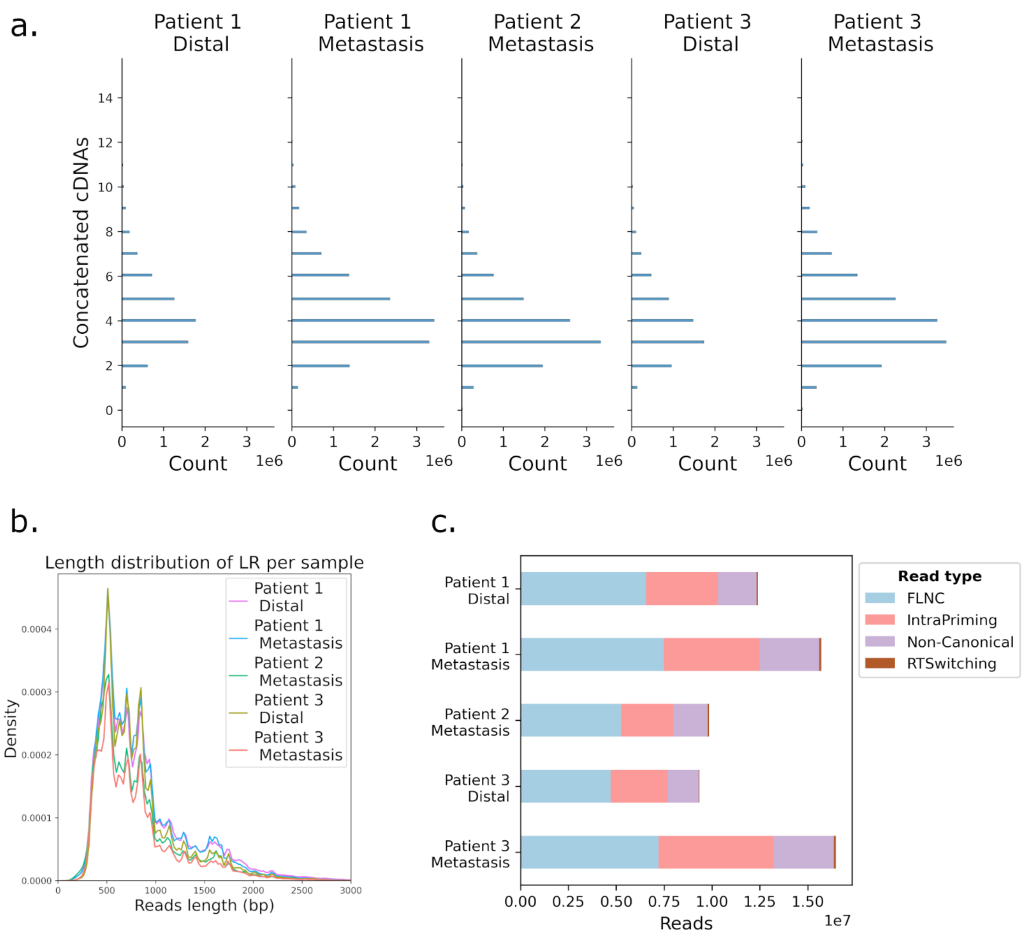


**Extended Data Figure 1: QC metrics long reads.** **(a)** Histogram showing the number of concatemeric cDNA molecules per sequencing read by sample. **(b)** Length distribution of long reads after unconcatenation, per sample. **(c)** Number of UMIs belonging to isoforms passing all filters (full-length non-chimeric) and UMIs belonging to filtered isoforms (intra-priming, reverse transcriptase (RT) switching, non-canonical isoforms), following SQANTI classification.

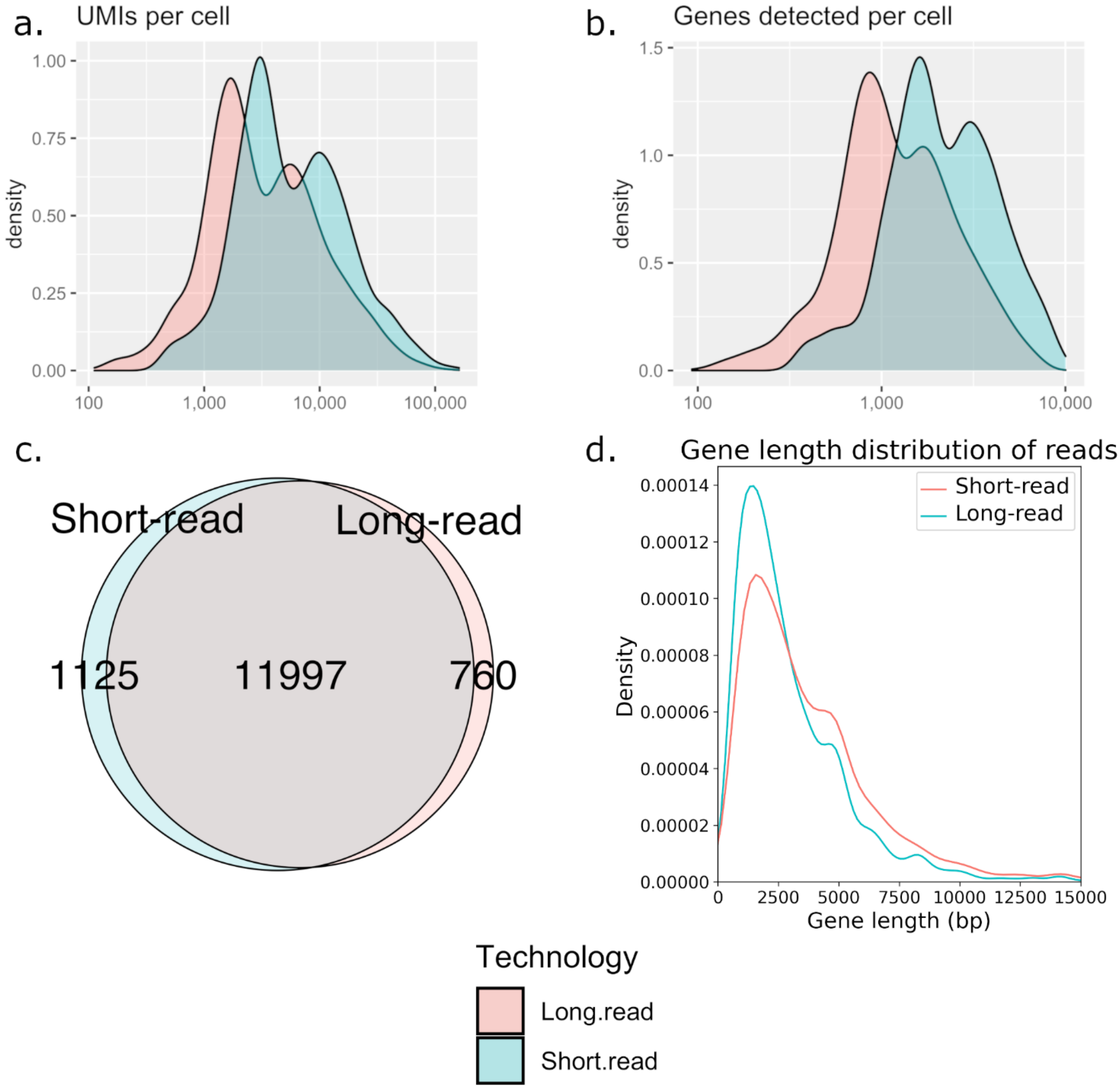


**Extended Data Figure 2: QC metrics comparison of short and long reads. (a)** Distribution of single UMI reads and **(b)** genes detected per cell in short- (light-blue) and long-read (light-red) technologies. **(c)** Overlap of total detected genes between short- and long-read datasets. **(d)** Gene length distribution of genes detected in short- and long-read technologies, where gene length equals the sum of exon length of the longest isoform associated with the gene.

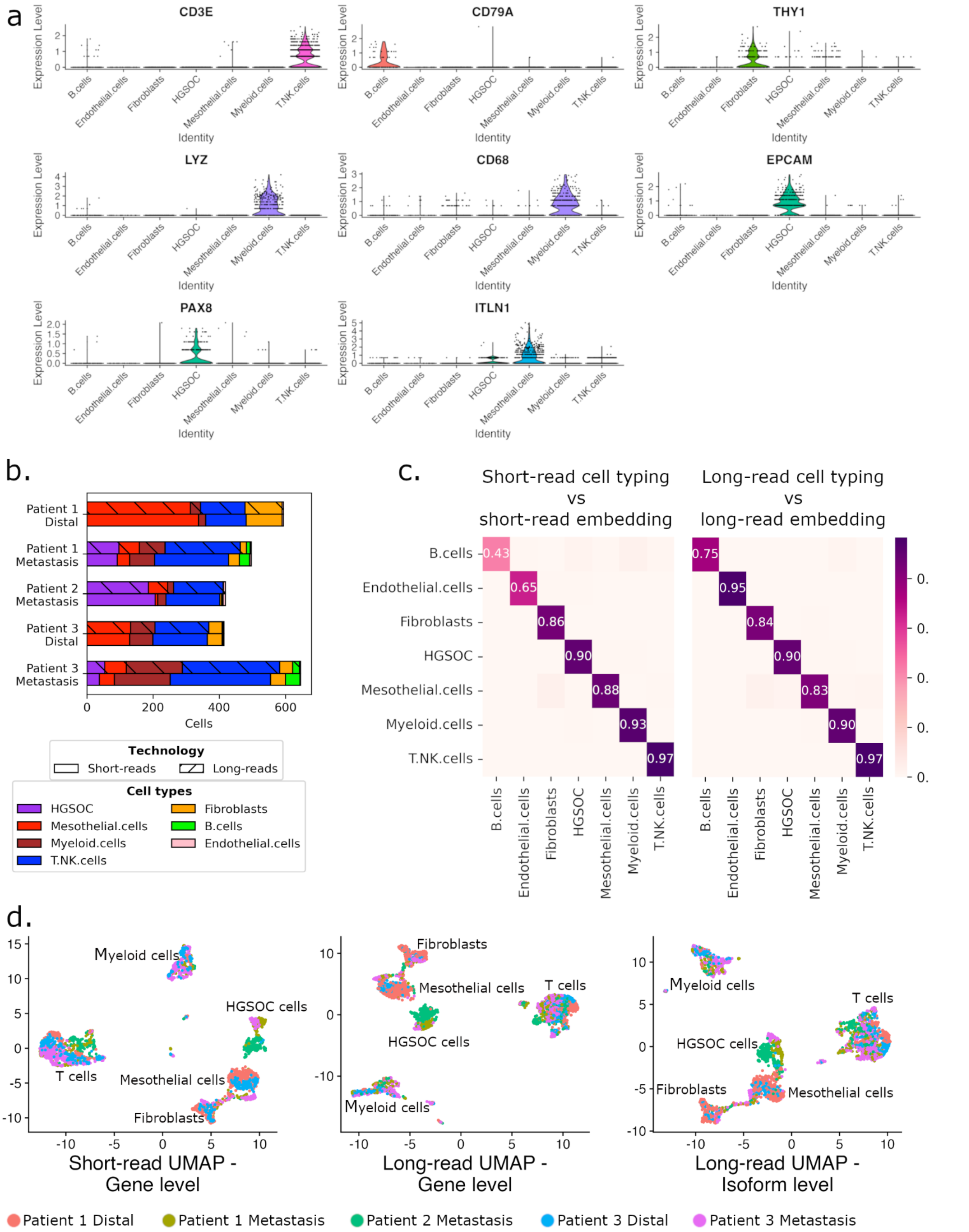


**Extended Data Figure 3: Cell type marker detection and embedding by sample. (a)** Violin plot showing expression of marker genes in long-read data. CD3E is a T cell marker, CD79A a B cell marker, THY1 a fibroblast marker, LYZ and CD68 myeloid cell markers, EPCAM and PAX8 are HGSOC markers and ITLN1 a mesothelial cell marker. **(b)** Cell type composition barplot per sample per sequencing technology. **(c)** Jaccard distance of cell populations in different UMAP embeddings: short-read cell labeling versus short-read UMAP embedding (left), and long-read cell labeling versus long-read UMAP embedding. **(d)** UMAP cohort visualization by sample ID for short-read - gene level (left), long-read - gene level (middle), and long-read - transcripts level (right) data.

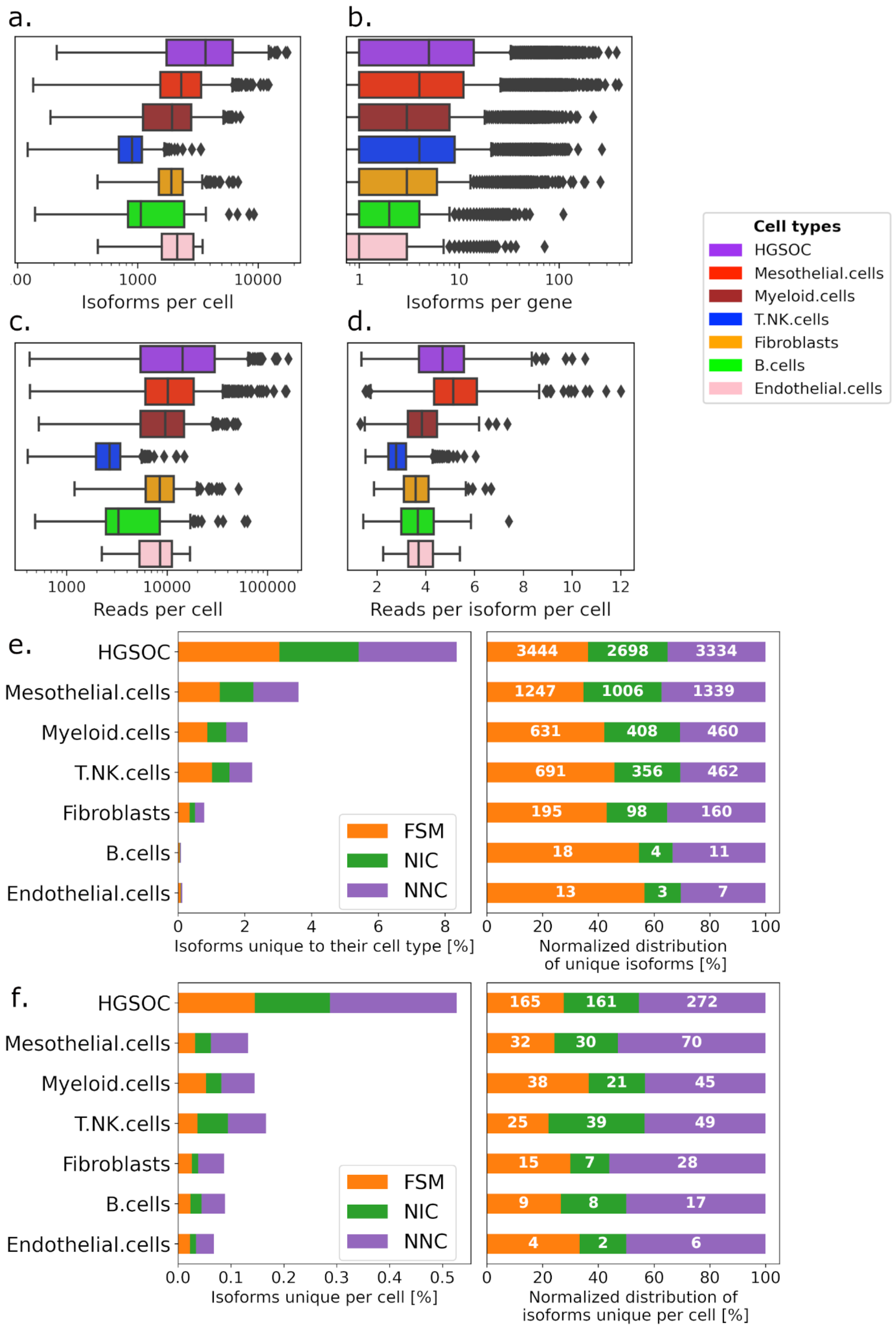


**Extended Data Figure 4: QC metrics by cell type in long-read sequencing. (a)** Number of isoforms detected per cell, by cell type. **(b)** Number of isoforms detected per gene, by cell type. **(c)** Number of unique reads per cell in long-reads, by cell type. **(d)** Long reads per isoform in each cell, by cell type. **(e)** Percentage of isoforms detected per cell type that are unique to this particular cell type (left panel), and their SQANTI-defined structural category normalized distribution (right panel, number of isoforms displayed in white). **(f)** Percentage of isoforms detected per cell type that are only found in one cell (left panel), and their SQANTI-defined structural category normalized distribution (right panel, number of isoforms displayed in white).

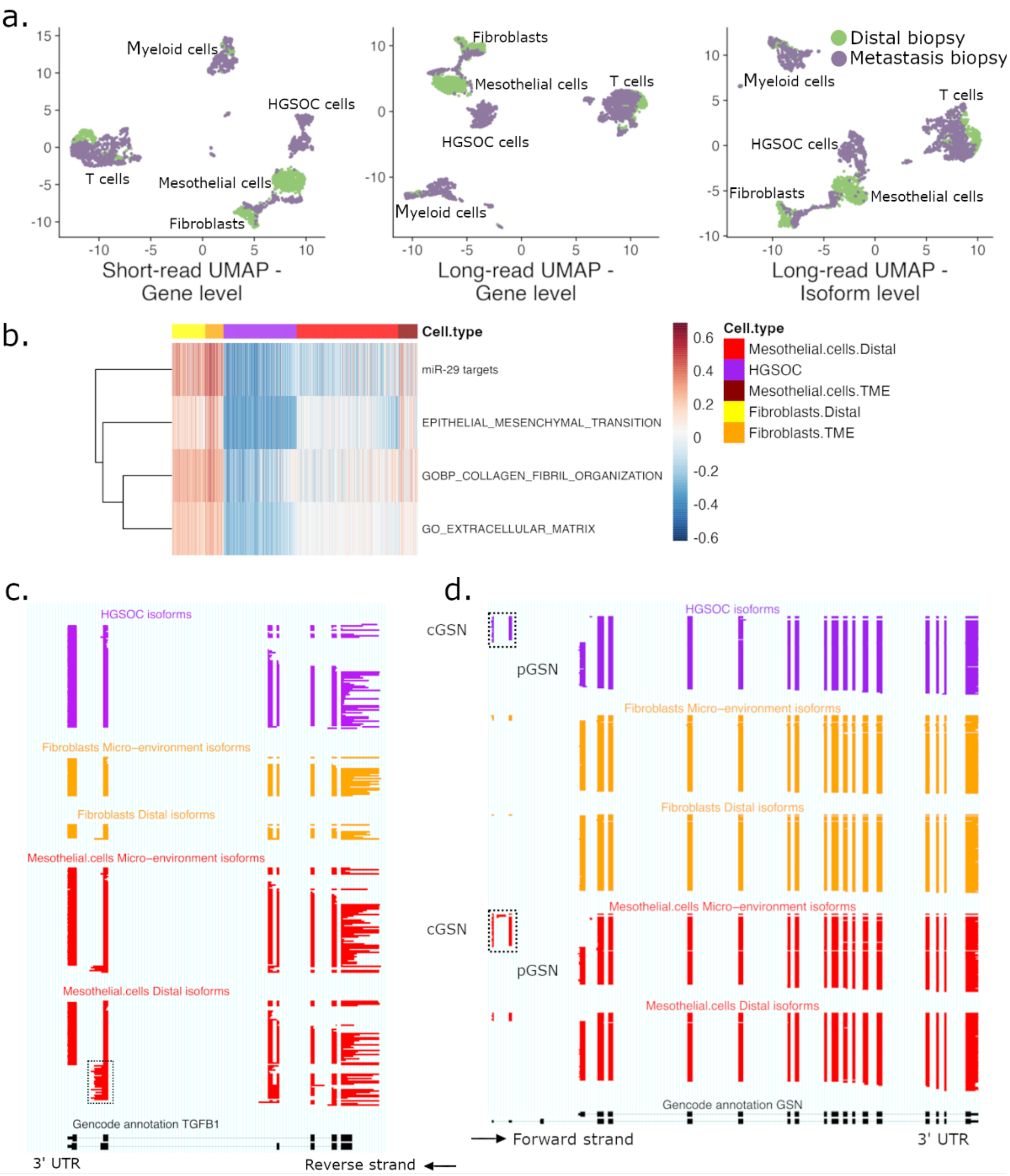


**Extended Data Figure 5: Epithelial-to-mesenchymal transition in the tumor microenvironment.** **(a)** UMAP embeddings of the cohorts' long-read data of short-read data - gene level (left), long-read data - gene level (middle), long-read data - isoform level (right), colored by tissue type. **(b)** Gene set variation analysis (GSVA) scores for different cell types. Heatmap colors from blue to red represent low to high enrichment. **(c)** ScisorWiz representation of isoforms in *TGFB1*. Dashed box highlights the non-canonical 3'UTR used by distal mesothelial cells. **(d)** ScisorWiz representation of isoforms in *GSN*. Dashed boxes highlight the TSS, where mesothelial TME and HGSOC cells differentially express the *cGSN* isoform, while mesothelial distal cells and fibroblasts use *pGSN*.

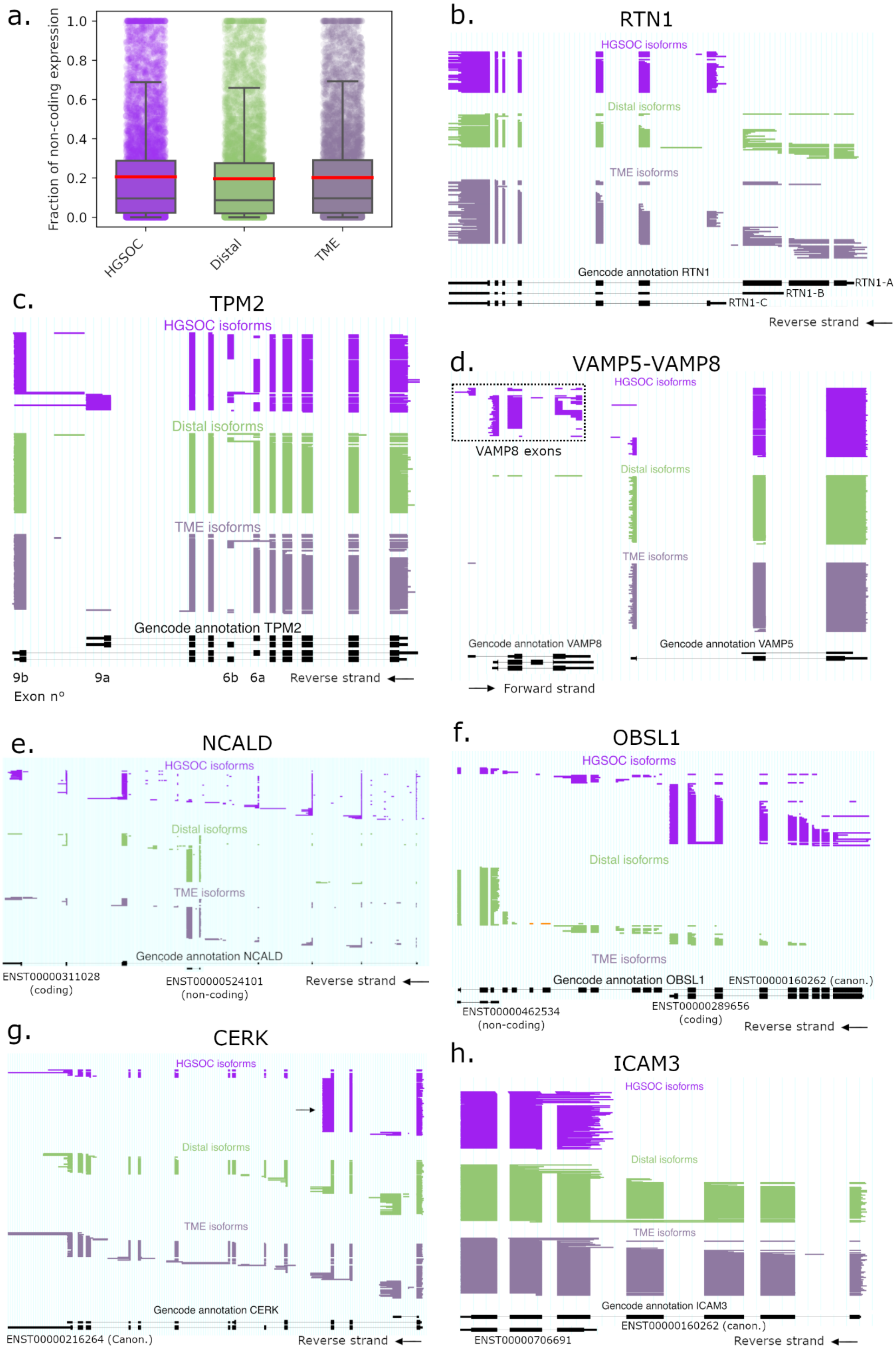
a.
Fraction of non-coding expression
1.0
0.8
0.6
0.4
0.2
0.0
HGSOC
Distal
TME
b.
RTN1
HGSOC isoforms
Distal isoforms
TME isoforms
Gencode annotation RTN1
RTN1-A
RTN1-B
RTN1-C
Reverse strand
c.
TPM2
HGSOC isoforms
Distal isoforms
TME isoforms
Gencode annotation TPM2
9b
9a
6b
6a
Reverse strand
Exon n°
d.
VAMP5-VAMP8
HGSOC isoforms
VAMP8 exons
Distal isoforms
TME isoforms
Gencode annotation VAMP8
Gencode annotation VAMP5
Forward strand
e.
NCALD
HGSOC isoforms
Distal isoforms
TME isoforms
Gencode annotation NCALD
ENST00000311028
(coding)
ENST00000524101
(non-coding)
Reverse strand
f.
OBSL1
HGSOC isoforms
Distal isoforms
TME isoforms
Gencode annotation OBSL1
ENST00000160262 (canon.)
ENST00000462534
(non-coding)
ENST00000289656
(coding)
Reverse strand
g.
CERK
HGSOC isoforms
Distal isoforms
TME isoforms
Gencode annotation CERK
ENST00000216264 (Canon.)
Reverse strand
h.
ICAM3
HGSOC isoforms
Distal isoforms
TME isoforms
Gencode annotation ICAM3
ENST00000160262 (canon.)
ENST00000706691
Reverse strand

**External Data Figure 6: Top differentially expressed isoforms between HGSOC and all distal cells.**

**(a)** Boxplot of the fraction of non-coding expression in protein-coding genes, where each point is a protein-coding gene**.** ScisorWiz representation of isoforms in **(b)** *RTN1*, **(c)** *TPM2*, **(d)** *VAMP5-VAMP8*, **(e)** *NCALD*, **(f)** *OBSL1*, **(g)** *CERK*, and **(h)** *ICAM3* in cancer, all distal, and all TME cells. Each horizontal line represents a single isoform colored according to cell types. Notable reference isoforms from GENCODE are named on the bottom of each gene.

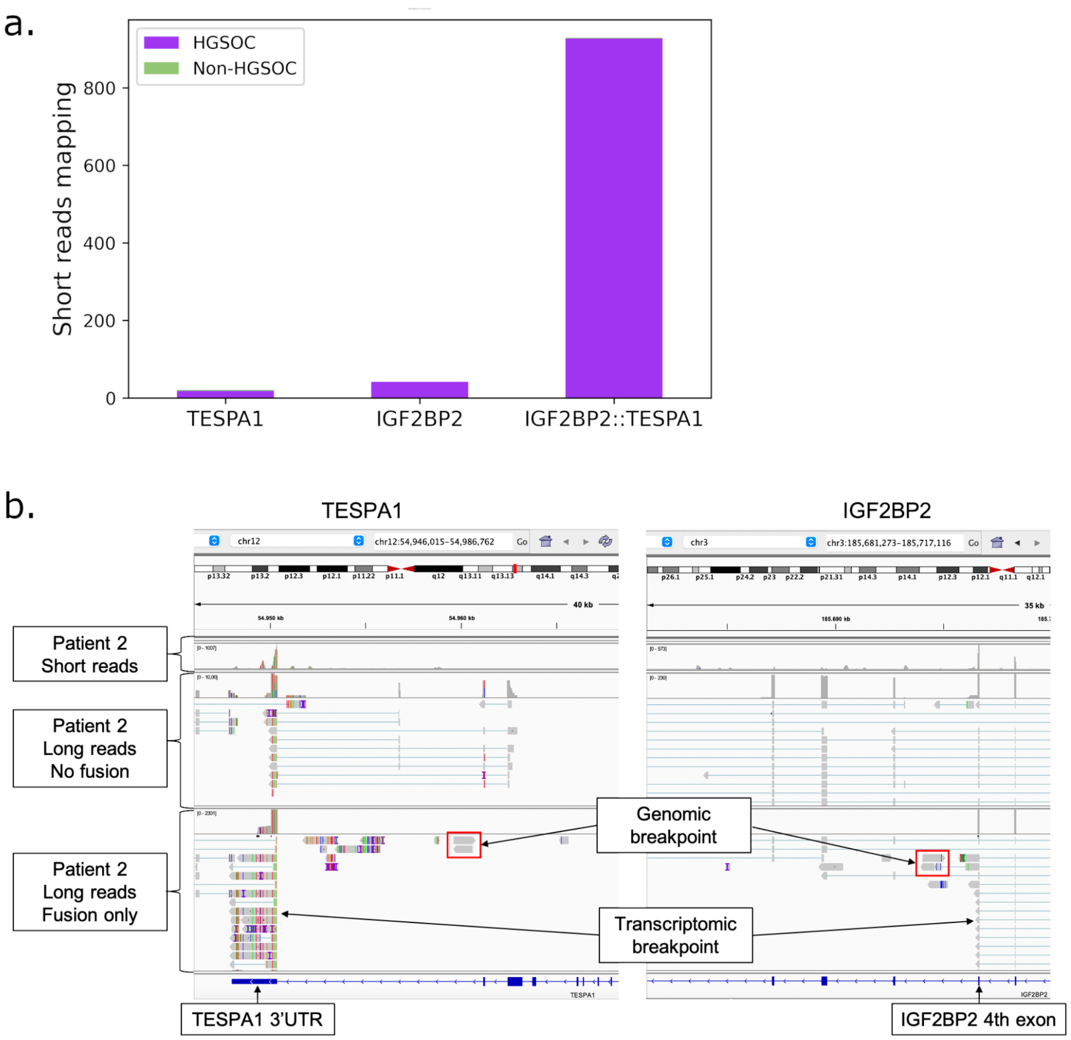


**Extended Data Figure 7: Transcriptomic and genomic breakpoint validation of the *IGF2BP2::TESPA1* gene fusion. (a)** Remapping of scRNA short-reads previously mapping to *IGF2BP2* or *TESPA1* and overlapping the fusion breakpoint. Most reads are preferentially remapped to the *IGF2BP2::TESPA1* transcript template. **(b)** Overview of *IGF2BP2::TESPA1* gene fusion genomic and transcriptomic breakpoints, on IGV genome browser. The two intronic fusion long reads in the red box share the same breakpoint and were used as template for the genomic fusion breakpoint reference. The transcriptomic breakpoint occurs on the closest 5' exon junction of the genomic breakpoint in *IGF2BP2*, and on the closest 3' exon junction in *TESPA1*.

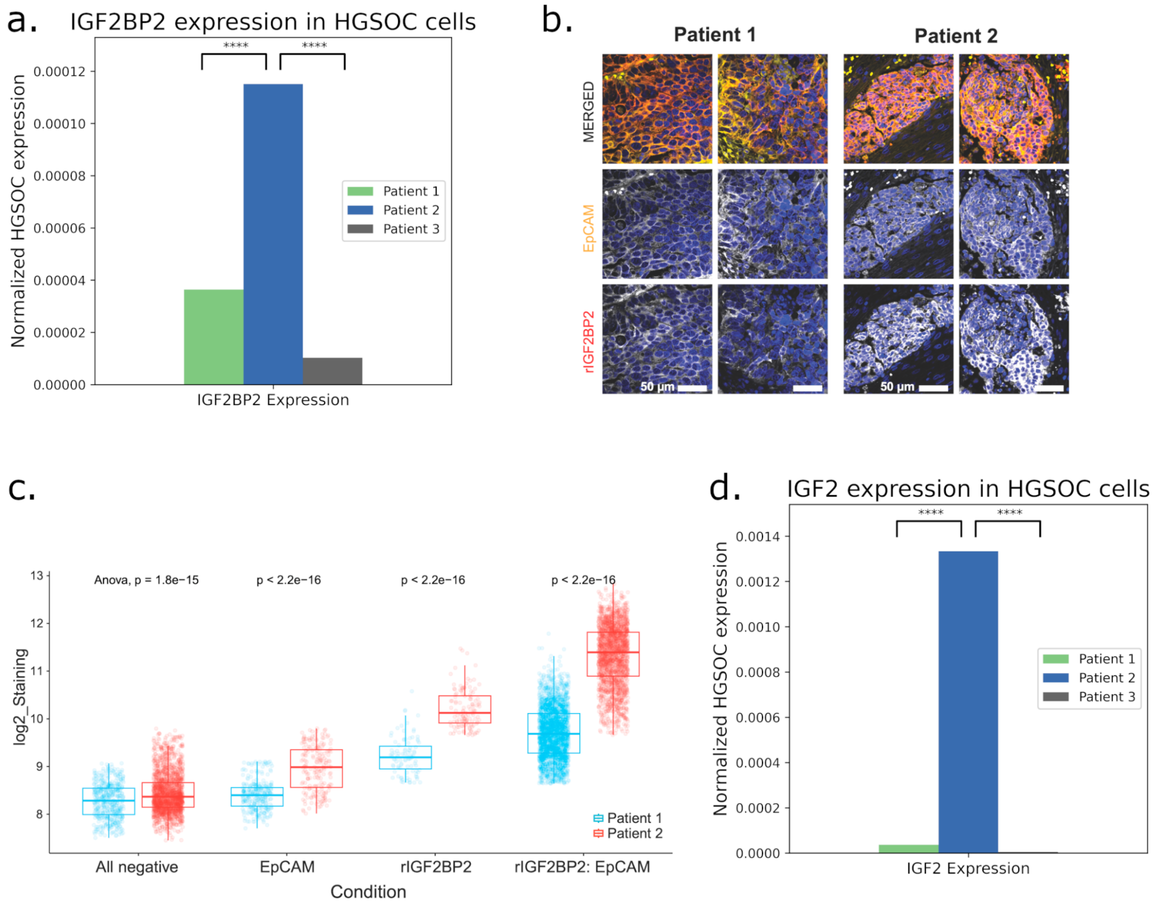


**Extended Data Figure 8: Immunofluorescence validation of IGF2BP2 expression.**
**(a)** Normalized expression of IGF2BP2 in cancer cells, per patient. **(c)** Representative immunofluorescence images (two tissue regions per patient) for detection of tumor (EpCAM$^{+}$) and C-terminal IGF2BP2 expressing cells in Patient 1 (no fusion) and Patient 2 harboring the *IGF2BP2::TESPA1* fusion. **(c)** Quantification of immunofluorescence images for determination of cancer cell-specific expression of IGF2BP2 in matched patient tissue samples. **(d)** Normalized expression of IGF2 in cancer cells, per patient

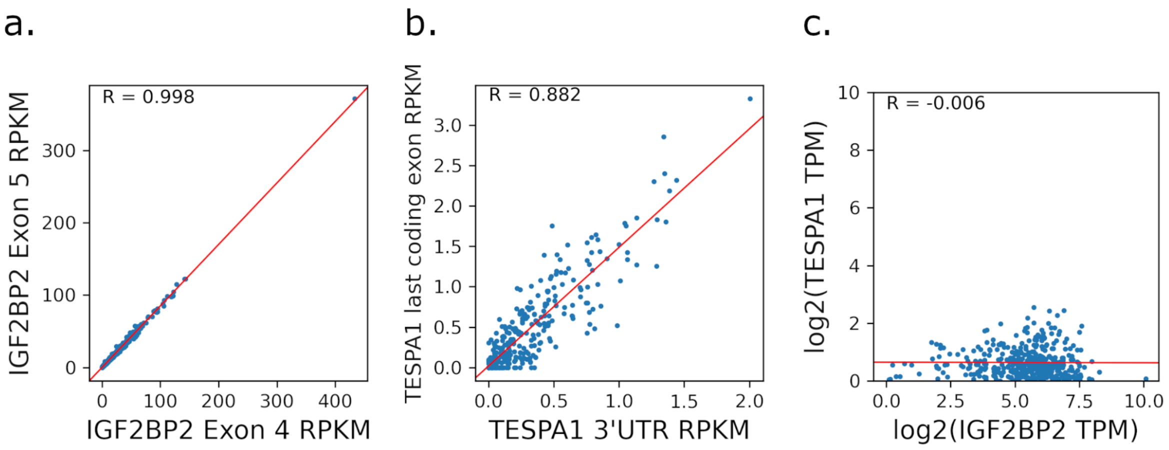


**Extended Data Figure 9**: **Detection of the *IGF2BP2::TESPA1* fusion in the ovarian cancer TCGA data.**

**(a)** TCGA expression of *IGF2BP2* exons 4 and 5, surrounding the *IGF2BP2::TESPA1* breakpoint. **(b)** TCGA expression of *TESPA1* last coding exon and 3'UTR exon, surrounding the *IGF2BP2::TESPA1* breakpoint. **(c)** Log2 normalized gene expressions of *IGF2BP2* and *TESPA1* in TCGA.

**Extended Data Table 1: Number of long reads per pre-processing step.** Full-length non-chimeric (FLNC) reads are the reads attached to isoforms passing all SQANTI filters.

| Patient | Biopsy site | #SMRT 8M cells | # cells | Sequencing output | Unconcatenated | Barcoded & poly-adenylated | Unique UMIs | FLNC | Av. reads/cell |
|---|---|---|---|---|---|---|---|---|---|
| Patient 1 | Distal omentum | 2 | 594 | 6,779,153 | 29,544,748 | 27,700,151 | 12,406,703 | 6,554,277 | 11034 |
| Patient 1 | Metastasis omentum | 4 | 497 | 13,384,311 | 57,548,787 | 53,882,454 | 15,705,384 | 7,033,927 | 14152 |
| Patient 2 | Metastasis omentum | 4 | 419 | 6,121,676 | 23,941,842 | 53,882,454 | 9,853,727 | 5,259,277 | 12551 |
| Patient 3 | Distal omentum | 2 | 415 | 14,110,642 | 58,681,499 | 22,312,230 | 9,358,572 | 4,698,768 | 11322 |
| Patient 3 | Metastasis omentum | 4 | 646 | 11,127,247 | 41,972,301 | 38,804,145 | 16,481,265 | 7,198,920 | 11143 |
| Total | / | 16 | 2571 | 51,523,029 | 211,689,177 | 196,581,434 | 63,805,651 | 30,745,169 | / |

**Extended Data Table 2: Number of short reads per pre-processing step.** scAmpi output includes the filtering of non-coding, mitochondrial and ribosomal genes.

| Patient | Biopsy site | # cells | Sequencing output | scAmpi output |
|---|---|---|---|---|
| Patient 1 | Distal omentum | 594 | 36,906,084 | 4,965,329 |
| Patient 1 | Metastasis omentum | 497 | 34,956,064 | 4,619,179 |
| Patient 2 | Metastasis omentum | 419 | 69,122,342 | 4,450,169 |
| Patient 3 | Distal omentum | 415 | 101,669,266 | 7,290,225 |
| Patient 3 | Metastasis omentum | 646 | 94,921,371 | 4,988,124 |
| Total | / | 2571 | 337,575,127 | 26,313,026 |
